# Chronic pain with use of analgesics and mortality in the home healthcare elderly: a nationwide population-based study

**DOI:** 10.1101/474239

**Authors:** Hsuan-En Chen, Wen-Ing Tsay, Shwu-Huey Her, Chung-Han Ho, Yi-Chen Chen, Kang-Ting Tsai, Chien-Chin Hsu, Jhi-Joung Wang, Chien-Cheng Huang

## Abstract

**Background:** Chronic pain may cause increased complications and all-cause mortality. However, nationwide data on elderly patients receiving home healthcare (HHC) remain unknown. Therefore, we conducted this study to address this issue.

**Methods:** We identified elderly individuals (≥ 65 years) with chronic pain receiving HCC between 2002 and 2013 in the Taiwan National Health Insurance Research Database. The comparisons of the causes of chronic pain, comorbidities, follow-up mortality, and the use of analgesics between two sexes and among three age subgroups were performed.

**Results:** A total of 1435 participants were identified, with a mean age of 77.8 ± 7.1 years and male percentage of 46.7%. The prevalence of chronic pain was 5.8%. Chronic pain was most prevalent in the 75–84 years age group (46.5%). Malignancy was the most common cause of chronic pain (94.2%), followed by peripheral vascular diseases (6.0%), osteoarthritis (4.3%), pressure ulcer (3.9%), spine diseases (3.1%), osteoporosis (1.3%), and headache (1.3%). The follow-up mortality was 32.8% within 6 month, 64.1% within 1 year, 79.9% within 2 years, and 84.3% within 3 years without difference in two sexes and age subgroups. Acetaminophen was found to be the most common analgesics, followed by non-steroidal anti-inflammatory drugs and opioids. Morphine was the most commonly used opioid.

**Conclusions:** This study delineates the causes of chronic pain, use of analgesics, and follow-up mortality in the HHC elderly, clarifying the relationship between chronic pain and the HCC elderly. This will facilitate the further investigation of this issue in the future.

## Introduction

Aging is a global issue in public health. In the United States, the geriatric population (≥ 65 years) is projected to be 20% of the total population by 2030, compared with 13% in 2010 and 9.8% in 1970 [1]. In the United Kingdom, the geriatric population was 18% of the total population in 2016 and projected to become 20.5% in 2026 and 23.9% in 2036 [2]. In Taiwan, the geriatric population was 13.3% in 2013, 14.0% in 2018, and will rapidly grow to 20% by 2025 [3,4]. Taiwan is one of the fastest aging countries in the world, therefore, the needs of long-term care services becomes a very urgent and important issue [5].

Home healthcare (HHC) services is one of the important components of long-term care [6]. It provides patients with continuous nursing care or physician visit needs following discharge from the hospital, through the provision of disease evaluations, nursing instructions, drug injections, fecal extraction, changing of urinary catheters or nasogastric tubes, or tube tracheostomy, nephrostomy or cystostomy catheters, changing the dressing of stage III and IV pressure sores, intravenous fluid injection, and ostomy nursing [6]. Chronic pain in the elderly is defined as “an unpleasant sensory and emotional experience associated with actual or potential tissue damage, or described in terms of such damage, for persons who are age ≥ 65 years with pain for greater than 3 months” [7]. Chronic pain may cause many complications in the elderly, including impaired activities of daily living, depression, deconditioning, polypharmacy, cognitive decline, poor quality of life, increased care burden and medical expenditure, and increased mortality [7-9]. Understanding the causes of chronic pain, its comorbidities, the use of medical or nursing resources, analgesic use, and subsequent mortality is crucial. However, we found nationwide data on this issue to be insufficient after an extensive search using the keywords “chronic pain”, “elderly”, and “home healthcare” via databases such as PubMed and Google Scholar. Therefore, we conducted this study to clarify this issue.

## Methods

### Data source

The Taiwan National Health Insurance Research Database (NHIRD) was used for this study. The NHIRD covers nearly 100% of the population’s healthcare data, and is one of the most comprehensive databases in the world [10]. The database contains registration files and original claim data for reimbursement and is provided to scientists for research purposes [10].

### Study design, setting, and participants

We conducted this nationwide population-based study by first identifying individuals receiving HHC between the years of 1997 and 2013 in the NHIRD (Figure 1). All geriatric HHC participants (≥ 65 years) receiving HHC before 2001, and lacking information on their sex and birthdays were excluded. Ultimately, 1435 HHC elderly with chronic pain were identified. Demographic variables including age, sex, causes of chronic pain, other comorbidities, iatrogenesis, living areas, use of medical resources including hospitalization and emergency department (ED) visit < 1 year, follow-up mortality, and use of analgesics were included in the analysis.

**Figure 1.**
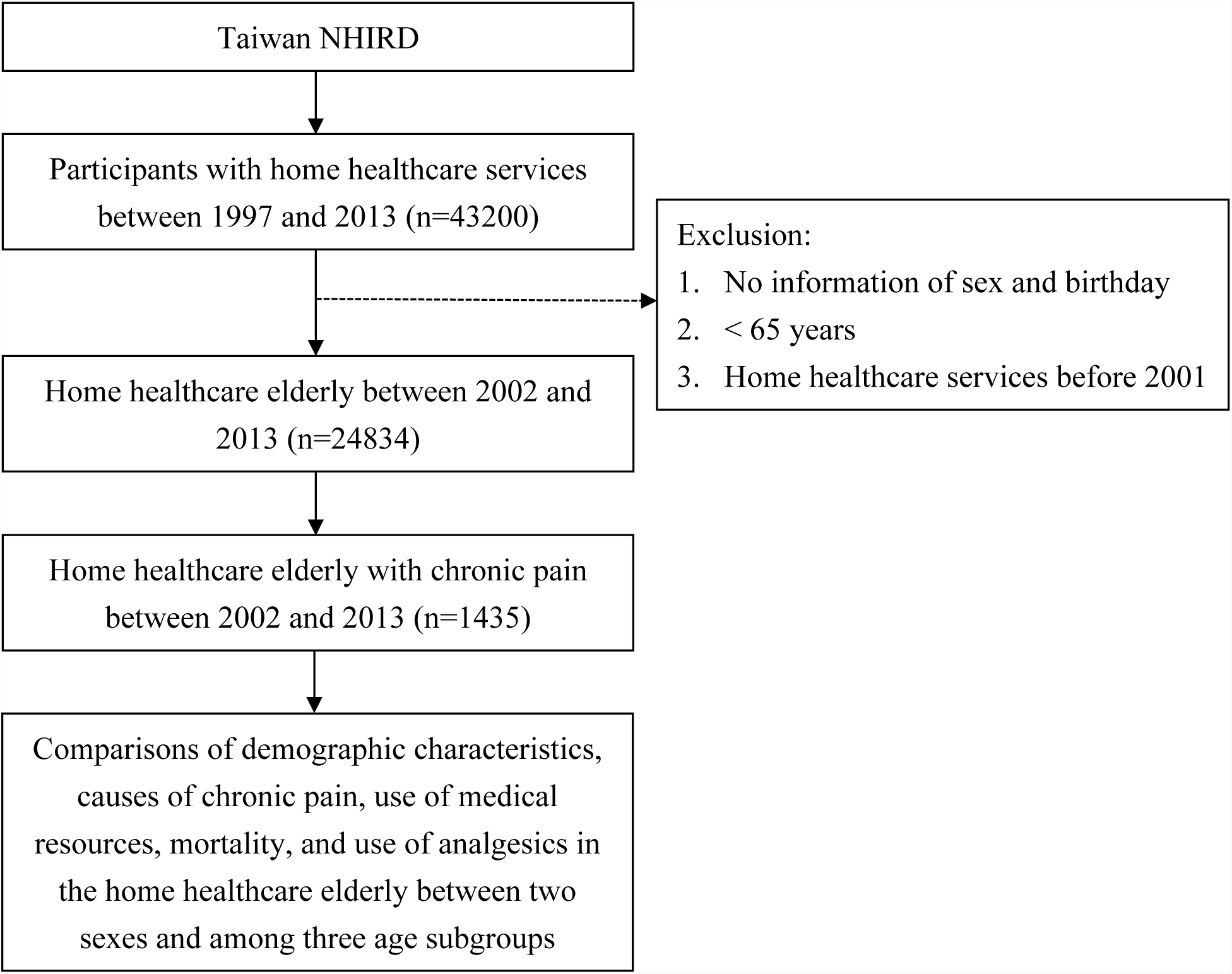
Flowchart of this study. NHIRD, National Health Insurance Research Database.

### Definitions of variables

Since the NHIRD does not provide a direct diagnosis of chronic pain, this study defines it as a characteristic of participants who have used either acetaminophen, non-steroidal anti-inflammatory drugs (NSAIDs; excluding aspirin), or opioids for at least 3 months, and also have been diagnosed with causes of chronic pain and comorbidities in at least one hospitalization or three out-patient clinics were defined as suffering from chronic pain.. The causes of chronic pain were defined as the follows: malignancy (International Classification of Diseases, Ninth Revision, Clinical Modification [ICD-9-CM]: 140-208), peripheral vascular diseases (ICD-9- CM: 443.8-444.9), osteoarthritis (ICD-9-CM: 715), pressure ulcer (ICD-9-CM: 707), spinal disorders (ICD-9-CM: 756.11, 756.12, 720-725, 737.1-737.4), osteoporosis (ICD-9-CM: 733.0), headache (ICD-9-CM: 307.81, 784.0, 346), herpes zoster (ICD-9-CM: 053), gout (ICD-9-CM: 274), rheumatoid arthritis (ICD-9-CM: 714), and diabetic neuropathy (ICD-9-CM: 250.6, 357.2). The other comorbidities were defined as follows: hypertension (ICD-9-CM: 401–405), diabetes (ICD-9-CM: 250), renal diseases (ICD-9-CM: 580-593), stroke (ICD-9-CM: 430-438), liver diseases (ICD-9- CM: 570-576), coronary artery disease (ICD-9-CM:410-414), chronic obstructive pulmonary disease (ICD-9-CM: 490-496), dementia (ICD-9-CM: 290, 291.2, 292.82, 294.1), hyperlipidemia (ICD-9-CM: 272), and depression (ICD-9-CM: 300.4). The age subgroups were classified as 65–74 years, 75–84 years, and ≥85 years [11-16].

### Ethical statements

This study was conducted in accordance with the Declaration of Helsinki and approved by the Institutional Review Board at Chi-Mei Medical Center. Since the data of the NHIRD have been stripped of personally identifiable details, informed consent was waived. The waiver does not affect the rights and welfare of the participants.

### Statistical methods

We used SAS 9.4 for Windows (SAS Institute, Cary, NC, USA) for all statistical analyses. Pearson chi-square tests were used for categorical variables, and independent t test or one-way ANOVA was used for continuous variables. The significance level was set at *p* < 0.05 (two-tailed).

## Results

A total of1435 HHC elderly with chronic pain were identified in this study (Figure 1 and Table 1). The mean age (± standard deviation) was 77.8 (± 7.1) years and the male percentage was 46.7%. The prevalence of chronic pain was 5.8% (1435/24834) in the HHC elderly. The age of 75–84 years was the largest age subgroup (46.5%), followed by 65–74 years (35.2%) and ≥ 85 years (18.3%). The common causes of chronic pain were malignancy (94.2%), peripheral vascular diseases (6.0%), osteoarthritis (4.3%), pressure ulcer (3.9%), spine diseases (3.1%), osteoporosis (1.3%), and headache (1.3%). In the comparison of causes of chronic pain between the two sexes, female participants had a higher percentage of osteoporosis (2.1% vs. 0.3%) than the male participants. In the comparison of other comorbidities, the male participants had higher prevalence rates of stroke and chronic obstructive pulmonary disease but lower prevalence rates of hypertension, diabetes, and hyperlipidemia than their female counterparts. Depression was diagnosed in only 0.4% of the total participants. Foley indwelling is the most common iatrogenesis (62.4%), followed by nasogastric tube (46.9%). More male participants had nasogastric tube and tracheotomy than the female participants. 36.9% the participants lived in South Taiwan, followed by those who lived in North Taiwan (35.3%). 76.7% of the participants were hospitalized and 52.8% of the participants visited the emergency department within 1 year after discharge from the hospital. The follow-up mortality was 32.8% within 6 month, 64.1% within 1 year, 79.9% within 2 years, and 84.3% within 3 years without difference between two sexes.

**Table 1.** Comparison of demographic characteristics, causes of chronic pain, use of medical resources, and mortality in the home healthcare elderly with chronic pain between two sexes.

| Variables | Total<br>n = 1435 | Male<br>n = 670 | Female<br>n = 765 | *p*-value |
| --- | --- | --- | --- | --- |
| Age (mean ± standard deviation) | 77.8 ± 7.1 | 77.6 ± 7.3 | 78.0 ± 6.9 | 0.322 |
| 65–74 years | 505 (35.2) | 217 (32.4) | 288 (37.7) | 0.115 |
| 75–84 years | 667 (46.5) | 325 (48.5) | 342 (44.7) |  |
| ≥ 85 years | 263 (18.3) | 128 (19.1) | 135 (17.7) |  |
| Causes of chronic pain* |  |  |  |  |
| Malignancy | 1351 (94.2) | 635 (94.8) | 716 (93.6) | 0.342 |
| Peripheral vascular diseases | 86 (6.0) | 46 (6.9) | 40 (5.2) | 0.193 |
| Osteoarthritis | 62 (4.3) | 30 (4.5) | 32 (4.2) | 0.784 |
| Pressure ulcer | 56 (3.9) | 25 (3.7) | 31 (4.1) | 0.754 |
| Spine diseases | 44 (3.1) | 23 (3.4) | 21 (2.8) | 0.451 |
| Osteoporosis | 18 (1.3) | 2 (0.3) | 16 (2.1) | 0.002 |
| Headache | 19 (1.3) | 10 (1.5) | 9 (1.2) | 0.601 |
| Herpes zoster | 13 (0.9) | 8 (1.2) | 5 (0.7) | 0.281 |
| Gout | 13 (0.9) | 9 (1.3) | 4 (0.5) | 0.102 |
| Rheumatoid arthritis | 10 (0.7) | 3 (0.5) | 7 (0.9) | 0.353 |
| Diabetic neuropathy | 6 (0.4) | 2 (0.3) | 4 (0.5) | 0.691 |
| Other comorbidities |  |  |  |  |
| Hypertension | 713 (49.7) | 313 (46.7) | 400 (52.3) | 0.035 |
| Diabetes | 378 (26.3) | 154 (23.0) | 224 (29.3) | 0.007 |
| Renal diseases | 265 (18.5) | 129 (19.3) | 136 (17.8) | 0.472 |
| Stroke | 246 (17.1) | 134 (20.0) | 112 (14.6) | 0.007 |
| Liver diseases | 231 (16.1) | 109 (16.3) | 122 (16.0) | 0.869 |
| Coronary artery disease | 184 (12.8) | 93 (13.9) | 91 (11.9) | 0.262 |
| Chronic obstructive pulmonary disease | 182 (12.7) | 125 (18.7) | 57 (7.5) | <0.001 |
| Dementia | 96 (6.7) | 42 (6.3) | 54 (7.1) | 0.550 |
| Hyperlipidemia | 88 (6.1) | 31 (4.6) | 57 (7.5) | 0.026 |
| Depression | 5 (0.4) | 3 (0.5) | 2 (0.3) | 0.669 |
| Iatrogenesis |  |  |  |  |
| Nasogastric tube | 673 (46.9) | 350 (52.2) | 323 (42.2) | <0.001 |
| Foley | 896 (62.4) | 409 (61.0) | 487 (63.7) | 0.307 |
| Tracheostomy | 100 (7.0) | 67 (10.0) | 33 (4.3) | <0.001 |
| Patient-controlled analgesia | 14 (1.0) | 1 (0.2) | 13 (1.7) | 0.003 |
| Living areas |  |  |  |  |
| North | 506 (35.3) | 233 (34.8) | 273 (35.7) | 0.116 |
| Center | 247 (17.2) | 113 (16.9) | 134 (17.5) |  |
| South | 530 (36.9) | 239 (35.7) | 291 (38.0) |  |
| East | 152 (10.6) | 85 (12.7) | 67 (8.8) |  |
| Use of medical resources < 1 year |  |  |  |  |
| Hospitalization | 1100 (76.7) | 538 (80.3) | 562 (73.5) | 0.002 |
| Emergency department | 757 (52.8) | 377 (56.3) | 380 (49.7) | 0.013 |
| Mortality | 1269 (88.4) | 603 (90.0) | 666 (87.1) | 0.082 |
| < 6 month | 471 (32.8) | 220 (32.8) | 251 (32.8) | 0.992 |
| < 1 year | 920 (64.1) | 439 (65.5) | 481 (62.9) | 0.297 |
| < 2 years | 1146 (79.9) | 543 (81.0) | 603 (78.8) | 0.295 |
| < 3 years | 1210 (84.3) | 572 (85.4) | 638 (83.4) | 0.305 |

Data were presented as number (percentage). \*Cause of chronic pain may be multiple.

The comparison of the causes of chronic pain among age subgroups showed that malignancy was more common in the 75–84 years group and peripheral vascular diseases was more common in the ≥ 85 years group (Table 2). The prevalence of hypertension, coronary artery disease, chronic obstructive pulmonary disease, and dementia increased with age. There was no difference of iatrogenesis, living areas, use of medical resources < 1 year, and follow-up mortality among three age subgroups.

**Table 2.** Comparison of demographic characteristics, causes of chronic pain, use of medical resources, and mortality in the home healthcare elderly with chronic pain among three age subgroups.

| Variables | Total | 65–74 years | 75–84 years | ≥ 85 years | *p*-value |
| --- | --- | --- | --- | --- | --- |
| Sex |  |  |  |  | 0.115 |
| Female | 765 (53.3) | 288 (57.0) | 342 (51.3) | 135 (51.3) |  |
| Male | 670 (46.7) | 217 (43.0) | 325 (48.7) | 128 (48.7) |  |
| Causes of chronic pain* |  |  |  |  |  |
| Malignancy | 1351 (94.2) | 478 (94.7) | 635 (95.2) | 238 (90.5) | 0.019 |
| Peripheral vascular diseases | 86 (6.0) | 26 (5.2) | 32 (4.8) | 28 (10.7) | 0.002 |
| Osteoarthritis | 62 (4.3) | 21 (4.2) | 30 (4.5) | 11 (4.2) | 0.954 |
| Pressure ulcer | 56 (3.9) | 18 (3.6) | 30 (4.5) | 8 (3.0) | 0.521 |
| Spine diseases | 44 (3.1) | 17 (3.4) | 22 (3.3) | 5 (1.9) | 0.478 |
| Osteoporosis | 18 (1.3) | 4 (0.8) | 10 (1.5) | 4 (1.5) | 0.510 |
| Headache | 19 (1.3) | 6 (1.2) | 11 (1.7) | 2 (0.8) | 0.535 |
| Herpes zoster | 13 (0.9) | 5 (1.0) | 5 (0.8) | 3 (1.1) | 0.826 |
| Gout | 13 (0.9) | 5 (1.0) | 6 (0.9) | 2 (0.8) | >0.999 |
| Rheumatoid arthritis | 10 (0.7) | 3 (0.6) | 7 (1.1) | 0 (0.0) | 0.277 |
| Diabetic neuropathy | 6 (0.4) | 3 (0.6) | 2 (0.3) | 1 (0.4) | 0.863 |
| Other comorbidities |  |  |  |  |  |
| Hypertension | 713 (49.7) | 224 (44.4) | 346 (51.9) | 143 (54.4) | 0.009 |
| Diabetes | 378 (26.3) | 140 (27.7) | 179 (26.8) | 59 (22.4) | 0.266 |
| Renal diseases | 265 (18.5) | 93 (18.4) | 133 (19.9) | 39 (14.8) | 0.195 |
| Stroke | 246 (17.1) | 84 (16.6) | 100 (15.0) | 62 (23.6) | 0.007 |
| Liver diseases | 231 (16.1) | 84 (16.6) | 106 (15.9) | 41 (15.6) | 0.915 |
| Coronary artery disease | 184 (12.8) | 45 (8.9) | 96 (14.4) | 43 (16.4) | 0.004 |
| Chronic obstructive pulmonary disease | 182 (12.7) | 39 (7.7) | 95 (14.2) | 48 (18.3) | <0.001 |
| Dementia | 96 (6.7) | 19 (3.8) | 44 (6.6) | 33 (12.6) | <0.001 |
| Hyperlipidemia | 88 (6.1) | 28 (5.5) | 43 (6.5) | 17 (6.5) | 0.791 |
| Depression | 5 (0.4) | 1 (0.2) | 3 (0.5) | 1 (0.4) | 0.852 |
| Iatrogenesis |  |  |  |  |  |
| Nasogastric tube | 673 (46.9) | 226 (44.8) | 321 (48.1) | 126 (47.9) | 0.486 |
| Foley | 896 (62.4) | 317 (62.8) | 408 (61.2) | 171 (65.0) | 0.541 |
| Tracheostomy | 100 (7.0) | 44 (8.7) | 40 (6.0) | 16 (6.1) | 0.161 |
| Patient-controlled analgesia | 14 (1.0) | 7 (1.4) | 5 (0.8) | 2 (0.8) | 0.529 |
| Living areas |  |  |  |  |  |
| North | 506 (35.3) | 166 (32.9) | 232 (34.8) | 108 (41.1) | 0.148 |
| Center | 247 (17.2) | 92 (18.2) | 122 (18.3) | 33 (12.6) |  |
| South | 530 (36.9) | 186 (36.8) | 250 (37.5) | 94 (35.7) |  |
| East | 152 (10.6) | 61 (12.1) | 63 (9.5) | 28 (10.7) |  |
| Use of medical resources < 1 year |  |  |  |  |  |
| Hospitalization | 1100 (76.7) | 397 (78.6) | 507 (76.0) | 196 (74.5) | 0.386 |
| Emergency department | 757 (52.8) | 263 (52.1) | 356 (53.4) | 138 (52.5) | 0.903 |
| Mortality | 1269 (88.4) | 452 (89.5) | 594 (89.1) | 223 (84.8) | 0.121 |
| < 6 month | 471 (32.8) | 163 (32.3) | 221 (33.1) | 87 (33.1) | 0.949 |
| < 1 year | 920 (64.1) | 337 (66.7) | 428 (64.2) | 155 (58.9) | 0.102 |
| < 2 years | 1146 (79.9) | 408 (80.8) | 536 (80.4) | 202 (76.8) | 0.387 |
| < 3 years | 1210 (84.3) | 428 (84.8) | 566 (84.9) | 216 (82.1) | 0.557 |

Data were presented as number (percentage). \*Cause of chronic pain may be multiple.

Acetaminophen was the most commonly used analgesic (86.5%), followed by NSAIDs (77.0%) and opioids (43.69%; Table 3). NSAIDs were more often used in the male participants. Morphine was the most commonly used opioid (27.3%). With advancing age, less participants used opioids as analgesics (65–74 years: 45.2% vs. 75–84 years: 44.2% vs. ≥85 years: 39.5%; Table 4).

**Table 3.** Comparison of analgesics in the home healthcare elderly with chronic pain between two sexes.

| Variable* | Total | Male | Female | *p*-value |
| --- | --- | --- | --- | --- |
| Acetaminophen | 1241 (86.5) | 589 (87.9) | 652 (85.2) | 0.138 |
| NSAIDs | 1105 (77.0) | 543 (81.0) | 562 (73.5) | <0.001 |
| Opioids | 627(43.69) | 275(41.04) | 352(46.01) | 0.058 |
| Morphine | 391 (27.3) | 174 (26.0) | 217 (28.4) | 0.309 |
| Fentanyl | 209 (14.6) | 86 (13.1) | 121 (15.8) | 0.151 |
| Pethidine | 27 (1.9) | 13 (1.9) | 14 (1.8) | 0.878 |

NSAIDs, nonsteroidal anti-inflammatory drugs. \*Use of analgesics may be multiple.

**Table 4.**
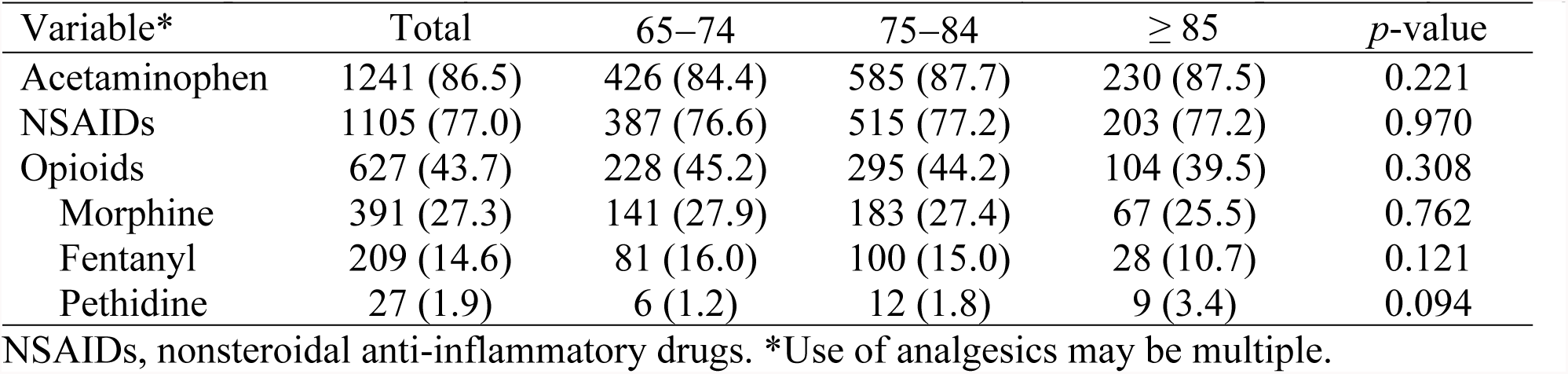
Comparison of analgesics in the home healthcare elderly with chronic pain among three age subgroups.

## Discussion

This study showed that the prevalence of chronic pain was 5.8% in the HHC elderly. Nearly half of the HHC with chronic pain belonged to the 75–84 years age group. Malignancy predominated as the cause of chronic pain and more common in the 75–84 years group than other age subgroups. Female participants were more likely to have osteoporosis as the cause of chronic pain than male participants. Although they received HHC services, 76.7% of the participants were hospitalized and 52.8% of the participants visited the ED within 1 year after discharge from the hospital. The followed-up mortality was high, 64.1% within 1 year, 79.9% within 2 years, and 84.3% within 3 years, without difference between two sexes and among three age subgroups. Acetaminophen was the most commonly used analgesic and morphine was the most commonly used opioid. There was a trend of decreased use of opioids with the age.

Contrary to common belief, the prevalence of 5.8% of chronic pain in the HHC elderly is surprising low. The possible explanation is the different criteria among studies. A convenience sample of community-dwelling elderly in Taiwan reported that the prevalence of pain was 50% [17]. Another study in Taiwan reported that the prevalence of pain in nursing home elderly was 65.3% [18]. However, both of the aforementioned studies utilized definitions of pain based on the self-reports of patients, which differs from the criteria for chronic pain in this study, since the strict criteria of “using of analgesics for at least 3 months” was also used. This additional criterion may have contributed to the low prevalence rate.

In this study, malignancy was the most common cause of chronic pain in the HHC elderly. The incidence and prevalence of most malignancies increase with the age [19]. Previous studies reported that 25-40% of geriatric patients with malignancy living at nursing home had daily pain [7,20]. However, 26% of these patients with daily pain received no analgesics and patients older than 85 years were more likely to receive no analgesia, which indicates that malignancy related pain may be undertreated in the geriatric population [20]. Regularly given oral analgesics are the most common treatment for the malignancy related pain in the elderly [21]. Because the elderly are at increased risk of developing toxicity from NSAIDs, opioids should be considered when the elderly have contraindications or uncontrolled pain after NSAIDs use [21]. The elderly need lower dose of opioids, therefore, careful titration based on individual response including the side effects can ensure the appropriate response to clinical demand [21]. In addition to analgesics, adjuvant medications including antidepressants, antiepileptics, corticosteroids and bisphosphonates may help in the treatment of certain types of chronic pain [21].

Females had more musculoskeletal pain and osteoporosis than males [22-24]. The prevalence of hip osteoporosis was 2% for men compared with 16% for women [24]. In women, the prevalence of osteoporosis rapidly increased (tripled) at the age of 70 years. In men, the prevalence of osteoporosis increased at the age of 80 years and the rate doubled [24]. In addition to the higher prevalence of osteoporosis, women may have higher pain sensitivity due to the more central sensitization [25].

The hospitalization and ED visit rate within 1 year after discharge were 76.7% and 52.8% in our study. It is difficult to compare previous data with this study because there is a great variation of medical care, insurance, culture, and local policy among different research. A study based on Medicare data reported that 20% of the beneficiaries (most are elderly) are rehospitalized within 30 days and 34% are rehospitalized within 90 days [26]. Another cross-national study reported that HHC elderly had a 25.3% of hospitalization and 15.7% of ED visit rates within 90 days after discharge [27]. The goals of HHC are to assist the patients to remain at home, avoid hospitalization or admission to long-term care institutions, and improve the patients’ function to give them greater independence [28]. A previous study revealed that an acute exacerbation of chronic disease is the most common reason for an unplanned hospitalization or ED visit, which can be prevented through knowledge of risk factors, provider communication, and careful monitoring [29]. Other risk factors are polypharmacy [30], the length of HHC episode [31], worsening primary or secondary diagnosis or the development of a new problem [32], falls [30], and advancing age [30]. Strategies recommended for preventing hospitalization or ED visit are patient/caregiver education, 24-hour on-call nursing coverage, telehealth, management support, case management, front loading visits, medication management, fall prevention program, and positive physician and hospital relationships [33].

The followed-up mortality in the HHC elderly with chronic pain was high in this study, however, there is also no comparable data in the literature. A study in Spain reported that 28.9% of HHC elderly died during the one-year follow-up period [34]. The independent predictors for morality were male sex, comorbidity, degree of pressure ulcers, and having received home care rehabilitation [34]. Another study about nursing home residents in Iceland reported that 28.8% of the residents died within a year, 43.4% within two years, and 53.1% of the residents died within 3 years [35]. The possible explanation for the higher mortality found in our study is that chronic pain and its underlying disease may increase mortality.

This study has the major strengths of possessing a nationwide design and presenting delineation of an unclear issue. However, there are also some limitations. First, we used “analgesics use for at least 3 months” as the surrogate criteria of chronic pain, which may be too strict and therefore result in an underestimation of the prevalence of chronic pain. Second, there may exist multiple causes of chronic pain, and the primary cause may be difficult to determine. Third, detailed information related to morality, including activities of daily living, life style, body-mass index, and socioeconomic status are not available in the NHIRD. Fourth, despite being a nationwide study, the results of this study may not applicable to other nations due to the differences in insurance, medical resources, race, and culture. Further studies are warranted to validate these result.

## Conclusions

This nationwide population-based study delineated that there was a prevalence of 5.8% of chronic pain in the HHC elderly in Taiwan. Malignancy was the most common cause. Chronic pain related to osteoporosis was more common in the female participants than in the male participants. The rates of hospitalization and ED visit were 76.7% and 52.8% within 1 year. The followed-up mortalities were 4.1% within 1 year, 79.9% within 2 years, and 84.3% within 3 years. Acetaminophen was the most commonly used analgesic and morphine was the most commonly used opioid. Further studies are warranted to validate these result.

## Abbreviations

HHC: Home healthcare
NHIRD: National Health Insurance Research Database
ED: emergency department
NSAIDs: Non-Steroidal Anti-Inflammatory Drugs
ICD-9-CM: International Classification of Diseases, Ninth Revision, Clinical Modification
ANOVA: Analysis of variance

## Declarations

### Consent for publication

Not applicable.

### Availability of data and materials

Data are available from the National Health Insurance Research Database (NHIRD) published by Taiwan National Health Insurance (NHI) Bureau. Due to legal restrictions imposed by the government of Taiwan in relation to the “Personal Information Protection Act”, data cannot be made publicly available. Requests for data can be sent as a formal proposal to the NHIRD (http://nhird.nhri.org.tw).

### Competing interests

The authors have declared that no competing interests exist.

### Funding

This study was supported by grant NHRINHIRD-99182 from the National Health Research Institutes in Taiwan and NSC 102-2314-B-384-001 from Taiwan National Science Council. The funders had no role in study design, data collection and analysis, decision to publish, or preparation of the manuscript.

### Authors’ Contributions

HEC, CC Huang, and JJW designed and conceived this study and wrote the manuscript. CHH and YCC performed the statistical analysis and wrote the manuscript. WIT, SHH, KTT, and CC Hsu provided professional suggestions and wrote the manuscript. All authors read and approved the final manuscript.

## Acknowledgments

We thank Miss Ti Hsu for the English revision.

